# EnsDTI-kinase: Web-server for Predicting Kinase-Inhibitor Interactions with Ensemble Computational Methods and Its Applications

**DOI:** 10.1101/2023.01.06.523052

**Authors:** Yijingxiu Lu, Sangsoo Lim, Sungjoon Park, MinGyu Choi, Changyun Cho, Soosung Kang, Sun Kim

**Author notes:** Equal contribution (Yijingxiu Lu, Sangsoo Lim).

## Abstract

**Motivation:** Kinase inhibitors are a major category of drugs. Experimental panel assay protocols are routinely used as a standard procedure to evaluate the efficiency and selectivity of a drug candidate to target kinase. However, current kinase panel assays are time-consuming and expensive. In addition, the panel assay protocols neither provide insights on binding sites nor allow experiments on mutated sequences or newly-characterized kinases. Existing virtual screening or docking simulation technologies require extensive computational resources, thus it is not practical to use them for the panel of kinases. With rapid advances in machine learning and deep learning technologies, a number of DTI tools have been developed over the years. However, these methods are yet to achieve prediction accuracies at the level of practical use. In addition, the performances of current DTI tools vary significantly depending on test sets. In this case, an ensemble model can be used to improve and stabilize DTI prediction accuracies.

**Results:** In this work, we propose an ensemble model, EnsDTI-kinase, that integrates eight existing machine learning and deep learning models into a unified model deployed as a web-server. Upon submission of a compound SMILES string, potential target kinases are automatically predicted and evaluated on the web-server. Importantly, EnsDTI-kinase is a computational platform where newly developed DTI tools can be easily incorporated without modifying core components so that its DTI prediction quality can improve over time. Besides, many useful functionalities are provided on our platform for users to further investigate predicted DTI: it allows confidence experiments by changing the amino acid (AA) at a specific position in a kinase sequence, named *in silico* mutagenesis, to investigate the effect of AA changes in binding affinity; it predicts kinase sequential regions where the query compound likely binds to by slidingly masking the sequence of selected kinases so that confidence in the predicted binding sites can be evaluated. Our model was evaluated in three experimental settings using four independent datasets, and showed accuracy of 0.82 compared to the average accuracy of 0.69 from five deep learning methods on the ChEMBL dataset. It achieved average selectivity of 0.95 within kinase families such as TK, CAMK and STE. For 8 out of 17 recent drugs, our model successfully predicted their interactions with 404 proteins at average accuracy of 0.82.

**Availability:** http://biohealth.snu.ac.kr/software/ensdti

## 1. Introduction

Protein kinases (PKs) play crucial roles in controlling a majority of signaling processes such as cell growth, differentiation, and apoptosis, which is often related to many human diseases [1, 2, 3]. Since the development of Imatinib as the first strategic small-molecule kinase inhibitor to treat chronic myelogenous leukemia (CML), small-molecule inhibitors are designed to target specific mutagenesis profiles on PKs. Examples of such inhibitors include erlotinib, sorafenib and gefitinib [4]. *In vitro* kinase assay has been widely used as a standard protocol of experimental evaluation for both efficiency and selectivity for the set of pre-defined PKs [5, 6]. However, the assay requires a lot of cost for testing different concentrations of an inhibitor, various inhibitor candidates or different sequence compositions of kinase proteins. Thus, machine learning (ML) based *in silico* drug-target interaction (DTI) prediction methods naturally gained interest as one of pre-screening strategies to narrow down the candidate PK targets for experimental validation [7, 8].

Deep learning methods demonstrated a good predictive power leveraging recent data accumulation by encoding data types such as strings, images, or graphs in many application areas [9, 10, 11]. Deep learning technologies have also been applied for DTI prediction [12, 13, 14]. However, deep learning-based DTI prediction methods are yet to achieve high prediction accuracy at the level of practical use and their performance highly depends on the selection of benchmark datasets in kinase inhibitor prediction (See Section Results). In this situation, ensemble methods have been successful to improve the overall prediction performance by integrating the base classifier results [15, 11]. EnsemKRR and EnsemDT are introduced to address the data imbalance problem by Ezzat et al. [16, 17] using decision tree and kernel ridge regression as base classifiers. DrugE-Rank combines six similarity-based methods to prioritize drugs by utilizing features and similarities of drugs and targets [18]. DeepFusionDTA generates a feature map from structure and sequence information via neural network and applies bagging-based lightGBM to predict binding affinity [19].

## 2. Motivation and Contribuitions

The widely used panel assays are laborious and time-consuming. Only a limited number of kinases can be tested in a single assay. Thus, powerful and accurate computational methods can be helpful to narrow down the list of kinases to be tested extensively. However, predictions of existing ML DTI methods are not at the level of such prescreening use. One critical limitation is that each method takes single encoding schemes of compounds and proteins as input. This constrains ML model to consider data only from a single point of view, which probably makes an upper bound on prediction performance and also limited power for generalizability. In addition, there is no DTI prediction tool that provides a web interface in which users can easily perform *in silico* experiments only with SMILES information before synthesizing the actual compound.

In this study, we present an ensemble method for DTI prediction for kinases, EnsDTI-kinase, where anyone can perform DTI prediction for kinases with SMILES information only. EnsDTI-kinase combines five deep learning models and three machine learning techniques to provide improved predictive power. Our contributions in this study are summarized as follows. First, EnsDTI-kinase allows a flexible combination of additional models by taking prediction outcomes from each of the base classifiers. In this architecture of EnsDTI, newly developed methods can be easily included over time so that DTI prediction accuracy can be improved always. Second, EnsDTI-kinase allows confidence experiments by changing the amino acid (AA) at a specific position in a kinase sequence, named *in silico* mutagenesis, to investigate the effect of AA changes in binding affinity. Third, EnsDTI-kinase predicts where the inhibitor binds to the kinase by sliding the masking sequences of selected PKs so that confidence in the predicted binding sites can be evaluated. We evaluated our model rigorously in three different performance evaluation settings to demonstrate the effectiveness of our ensemble strategy. EnsDTI-kinase is publicly available at http://biohealth.snu.ac.kr/software/ensdti.

## 3. Materials and Methods

### 3.1. Data

#### 3.1.1. Datasets

We used four different DTI datasets: davis, kiba, kinome, and human. Below are brief descriptions of each of the datasets. The data sizes and characteristics of each dataset are summarized in Table 1.

**Table 1:** The summary of DTI databases used in this study. Detailed information on data pre-processing are described in Section 3.1.

| **dataset** | **DTIs** | **P/N ratio** | **Compounds** | **Kinases** |
| --- | --- | --- | --- | --- |
| davis | 28,288 | 1:11 | 64 | 442 |
| kiba | 118,254 | 4:1 | 2,111 | 229 |
| kinome | 48,547 | 1:1.6 | 28,860 | 346 |
| human | 6,212 | 1.2:1 | 1,052 | 2,504 |

*davis* dataset [20] is a well-known benchmark dataset for binding affinity prediction of kinase inhibitors. It includes experimentally determined dissociation constant (*K*_*d*_) values of 68 kinase inhibitors against 442 kinase proteins, resulting in a total of 30,056 DTIs.

*kiba* dataset is a kinase-inhibitor bioactivity dataset [21] that integrated a total of 246,088 DTIs for 52,498 chemical compounds and 467 kinases. In this study, we used a reduced version of the kiba dataset retrieved from He et al. [22] to make sure that each target kinase has at least 10 DTIs. The filtered kiba dataset contains 118,254 DTIs, 229 kinases and 2,111 compounds.

*kinome* dataset is used in MDeePred [23]. The dataset is originally collected from ChEMBL (version: 25) on selected compounds from clusters by ECFP4 fingerprints with Tanimoto similarity over 0.7. The final dataset consists of 346 proteins, 28,860 compounds, and 48,547 DTIs.

*human* dataset was used in Tsubaki et al. [24], originally curated by Liu et al. [25]. The dataset contains 3,364 positively labeled DTIs on 1,052 compounds and 2,504 proteins. According to the authors, dissimilar proteins are less likely to be targeted by similar ligands. Based on the assumption, they screened 2,848 highly credible negative DTI pairs.

#### 3.1.2. Data Pre-processing

We removed DTIs whose compounds contain any non-covalent bonds. This resulted in 64, 2111, 28860 and 1052 compounds for davis, kiba, kinome and human datasets, respectively. For datasets that provide continuous affinity values, we binarized them so that data can be used for a classification problem. Specifically, we used *pK*_*d*_ of 7.0 and 12.1 as binarization cutoff values for davis and kiba datasets, respectively, as suggested by Tang et al. [21]. Detailed information (e.g. the positive-negative class label ratio, and the number of samples) about the datasets are in the Table 1.

### 3.2. Encoding

We used eight base classifiers (five deep learning and three machine learning models) in this study (See Section 3.3). Among these base classifiers, DL models have their own data encoding schemes while ML models lack benchmark encoding schemes. Thus, we chose data encoding schemes for the three ML models independently.

For DL models, compound information is used in SMILES format for CPI-Prediction [24], DeepDTA [26], and DeepPurpose [27], while as fingerprints for MDeePred [23] and DeepConv-DTI [28]. Protein information is used for DeepDTA, DeepConvDTI and DeepPurpose as AA sequence, for CPI-Prediction as 3-gram AA sequence, and for MDeePred as five fixed-size feature matrices. For ML models, compounds were encoded by Morgan fingerprints (256 bits) for all three ML models. We used three different protein descriptors considering structural and physico-chemical characteristics: AA composition (AAC) [29], pseudo-AA composition (PAAC) [30], and dipeptide deviation from expected mean (DDE) [31] for MLP, SVM and RF models, respectively. AAC is a 20-dimensional vector to reflect the occurrence frequency of 20 native AAs in a sequence. PAAC introduces a set of discrete numbers to approximate the effect of AA sequence order to generate an 80-dimensional vector. It combines both AA components and different sequence correlation factors while maintaining the same mathematical framework as AAC. DDE is an amino acid composition-based feature vector differentiating exact epitopes from non-epitopes. DDE is known to be able to capture both the relative positional information in a protein sequence and evolutionary information when detecting DNA-binding proteins.

Both `RDKit` [32] and `iFeature` [33] libraries were used to generate the aforementioned chemical and protein representations.

### 3.3. Model Architecture

The basic idea of EnsDTI-kinase is that data encoding, model architecture, and final prediction modules can be independently designed and built as separate components in a pipeline so that newly developed DTI models can be easily combined to the *training* module and independently participate in ensemble prediction for future improvements. In EnsDTI-kinase, we integrated eight base classifiers: five DL models such as DeepDTA [26], DeepConv-DTI [28], CPI-Prediction [24], DeepPurpose [27] and MDeePred [23]; three ML models such as SVM, RF, and MLP. Our ensemble model consists of three parts: *encoding, training* and *ensemble* (Figure 1). The *encoding* module transforms both compound and protein data into computable formats (See Section 3.2). The *training* module consists of the eight base classifiers where the models are independent of each other in training the encoded data. The classifiers were trained and evaluated using the datasets prepared in this study (Details in Section 3.3.1). The *ensemble* module combines predictions from the *training* module and generates the final prediction using a series of fully connected layers. In the ensemble module, we combined the eight pairs of positive and negative probability values from the eight base classifiers to make the final prediction. Instead of using a voting scheme, we trained a fully-connected layer by applying GridSearchCV of `scikit-learn` package [34] to exhaustively search model parameters through a number of hidden neurons among {3, 4, 5}.

**Figure 1:**
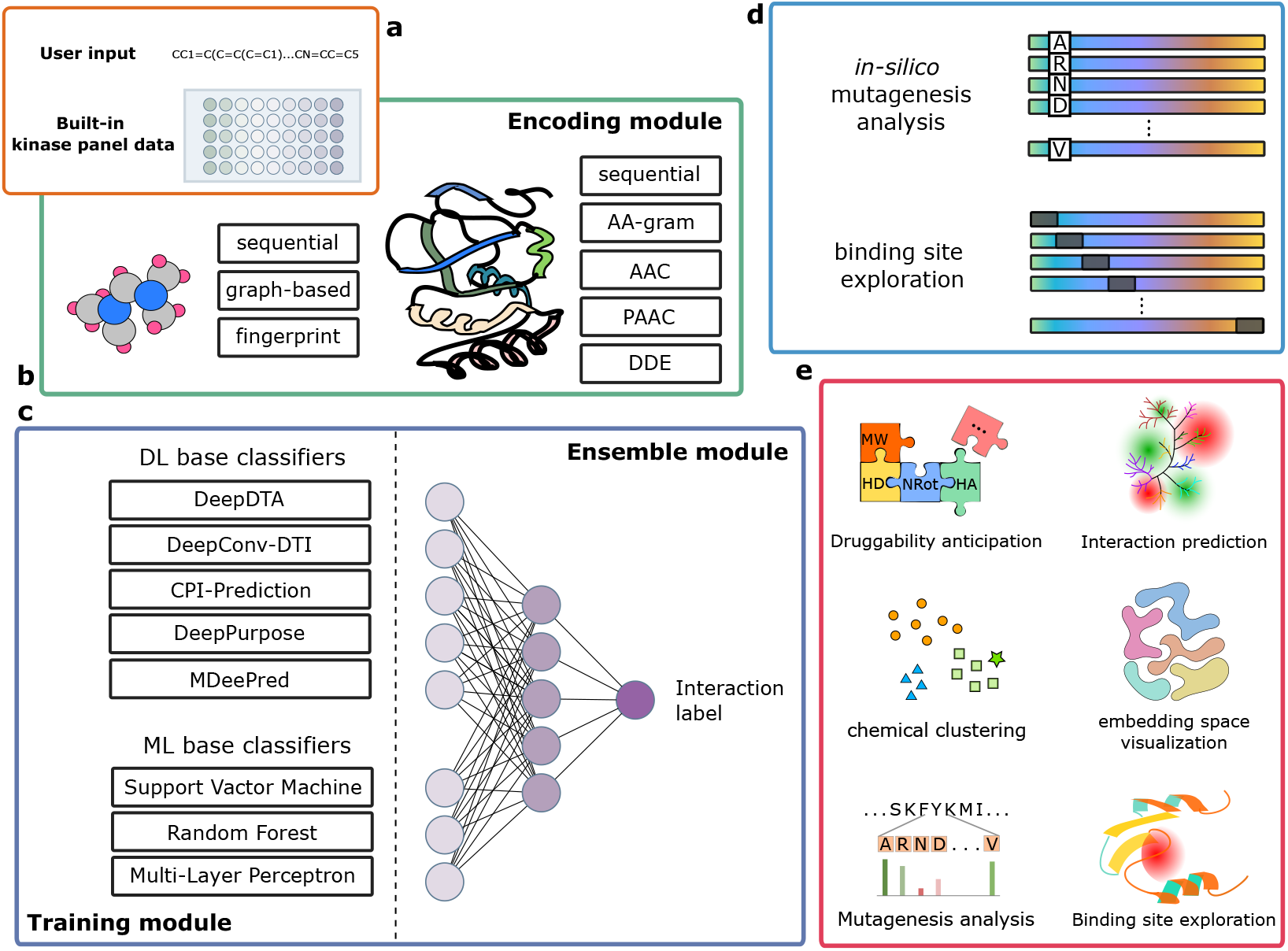
Framework of EnsDTI-kinase: **(a)** Given a SMILES string of query compound and the built-in kinase panel data as input, we first **(b)** encode both proteins and compound into model-learnable vectors considering their structural and functional characteristics (**encoding** module). **(c)** Then, we send these vectors into our ensemble framework to predict the interaction label of each pair of compound and protein (with **training** and **ensemble** module). **(d)** We also design two components, *in silico* mutagenesis analysis and binding site exploration, to provide model explainability. **(e)** Both of the two components are implemented with useful functions in our web-server.

**Figure 2:**
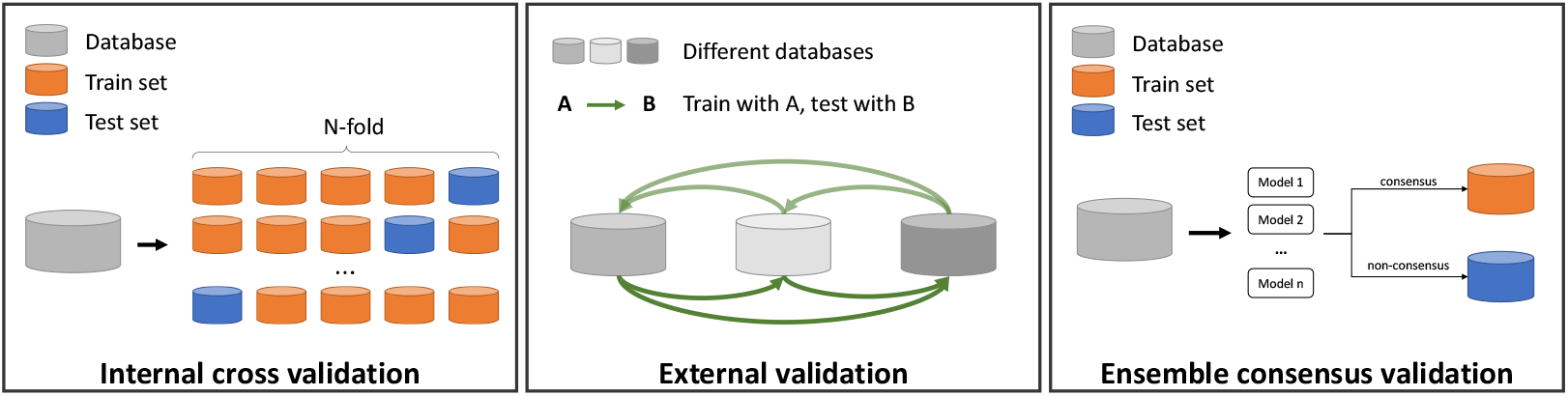
Three validation schemes used in this study: a) Internal cross-validation: randomly split each dataset into 5 folds and use the 3 folds for training, one of the other folds for validating, and the remaining fold for testing in turn. b) External validation: given several databases from different sources, use each of them for training and validating, and each of the rest databases for testing. c) Ensemble consensus validation: for davis dataset, we split the database into train and test datasets according to the consensus results of credible SOTA deep-learning models.

#### 3.3.1. Base Classifiers

##### Deep Learning Classifiers

We chose five DL classifiers: MDeePred [23], CPI-prediction [24], DeepDTA [26], DeepPurpose [27], and DeepConv-DTI [28]. These methods generate latent patterns of protein and compound separately and combine them to make the final prediction. The output size for all the models was set to two: positive and negative binding probabilities. Detailed information on each model will be explained in the following paragraphs.

DeepDTA and MDeePred were designed for regression problems. For the classification task, we modified the activation and loss functions and model parameter settings slightly so that the model could be applied to classification task. To be specific, we change the activation function for the regression task to a sigmoid function for the binary classification prediction 1 and calculate loss with binary cross entropy 2. The output size is set to two to represent positive and negative binding probabilities.

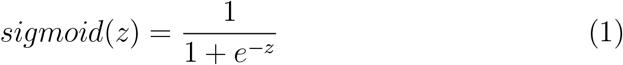

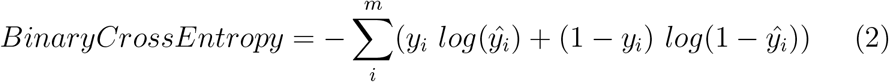

##### DeepDTA

DeepDTA [26] uses sequential information for proteins and compounds. By representing each character of SMILES strings and amino acid in one-hot encoding, DeepDTA uses two blocks of convolution neural networks (CNNs) to capture compound and protein features separately, followed by a concatenation of both representations for final prediction through an MLP layer.

##### DeepConv-DTI

DeepConv-DTI [28] extracts latent information separately for protein and compound. It employs CNNs on various lengths of protein subsequences and a fully connected layer (FC layer) on Morgan fingerprints to capture the local sequential patterns of DTI.

##### CPI-Prediction

CPI-prediction [24] learns chemical features from 2D molecular graphs and protein sequences with GCN and CNN, respectively. Protein sequences are encoded as a sequence of words that consist of 3-gram amino acid fed to 1D-CNN for latent representation. An attention mechanism is then used to calculate the relationship between each compound and these 3-gram amino acids.

##### DeepPurpose

DeepPurpose is an encoder-decoder framework-based model for DTI prediction [27]. It learns two embeddings for SMILES and AA sequences and feeds them into a CNN decoder to predict DTI binding affinity. Among various options they provide to build architecture, we chose message passing neural network (MPNN) to obtain the latent vector of a molecular graph and CNN to encode protein features from decomposing target sequence for binding AAs finding. Both features are sent to the decoder for DTI prediction.

##### MdeePred

MDeePred [23] uses protein encoding matrix and four different AA feature matrices, such as amino acid substitution matrix, physicochemical property difference matrix, distance-dependent statistical potential matrix, and residue contact energy matrix, to represent protein. A multi-channel CNNs was used to combine these 5 features. Compounds are represented as fingerprints and fed to an independent feed-forward network in parallel.

##### Three ML Base Classifiers

We implemented ML base classifiers using `sklearn` Python package. According to the parameter settings for MLP, the number of hidden layers is set to one and the number of neurons is 100. We chose ReLU as the activation function, and learning rate of 1e-3. We implemented SVM with regularization parameter equal to 1.0, with radial basis function (RBF). We implemented RF with the number of trees equal to 100, and the Gini impurity as the function to measure the quality of split.

### 3.4. Evaluation

To rigorously evaluate the performance of EnsDTI, we designed three different experimental settings.

#### Internal Cross-Validation

For each dataset in Table 1, DTI data were divided into 5-fold to build training, validation, and testing sets with a ratio of 3:1:1.

#### External Validation

As we have prepared four independent datasets from different sources, EnsDTI-kinase was evaluated by training an ensemble model on one of the datasets and using the model for prediction on the other independent dataset. This experiment evaluates how much EnsDTI-kinase can be generalized to previously unseen data.

#### Ensemble Consensus Validation

We next conducted an experiment to resample data by considering two aspects from a machine-learning perspective. First, we redefined the train and test sets by majority voting of the five DL models with threshold of 3. DTIs classified by three or more DL models are considered as training data and the rest as test sets. In other words, we can evaluate how well the classifiers trained on easily predictable samples (i.e. distant from decision boundary) can predict difficult samples (close to decision boundary). Second, as most DTI datasets contain much more negatively-labeled data points than positively-labeled ones, the five DL models were high in consensus prediction on the negative samples. We removed negatively-labeled DTIs that were correctly predicted by all models from training set to prevent the model from being negatively biased. Our intention is to remove examples that any of models can easily predict negatively. Inclusion of such easy examples guides ML models to set decision boundary between positive and negative samples is not as tight as it should be. We sampled training data class balanced. As a result, we obtained 594 and 1,912 positives and negatives in training data, while 1,699 positives and 862 negatives in test sets.

### 3.5. Experiments for Prediction Confidence Evaluation

To provide explainability on our model, we introduce two components below:

- *in silico point mutagenesis -* investigation of the effect of sequential variation on binding affinity by *silico* mutagenesis to specific AAs of target kinases.
- *binding site prediction -* prediction of the potential binding sites by sliding a window of sequential perturbation.

We employed the perturbation-based interpretation method [35] to build an AA sequence-wise importance map to investigate the contribution of each residue in the given protein. We collected 33 and 11 structured protein-compound pairs from Roskoski Jr [36] and Wu et al. [37] to interpret our model, respectively. We generated a series of masked sequences for each protein sequence by sliding window. These masked sequences are fed into our model with the same SMILES string to calculate the perturbed interaction value. Besides, we calculate the distance between inhibitor and kinase residues according to their crystal structure. These structure-based distance values are used as reference for perturbation analysis. By comparing the reference and the perturbation values, we can evaluate to what extent can our model extract potential pocket information from the given kinase and inhibitor pairs.

Specifically, perturbed interaction value is calculated by comparing the occlusion result and the original predicted value through the function presented in Equation 3. The measure of masked region importance *K*_*ij*_ is defined as the division of the masked prediction deviation by the prediction error, where *v*_*i*_ represents the actual binding label of protein *i*, and predicted value before and after occlusion is represented as *p*_*i*_ and *p*_*ij*_ respectively.

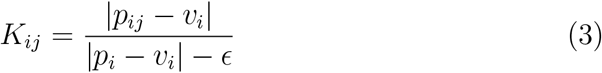

One major hurdle for displaying interpretation results is that majority of panel kinases have not been crystallized. Therefore, many protein binding site prediction models showed a limited application focusing on the heatmap generation, rather than projecting on protein structures. Fortunately, all kinase structures could be obtained by using AlphaFold2. Google Deepmind recently released an [AlphaFold Protein Structure Database](https://alphafold.ebi.ac.uk/) which contains the human proteome and majority of manually curated UniProt entries [38, 39]. Interpretation results and motif annotation references from the kinase-ligand interaction fingerprints and structure database (KLIFS) were both projected on these numerically predicted kinase structures.

### 3.6. Web-server Implementation

EnsDTI-kinase is implemented using Django 2.0 framework and SQLite 3 database for persistent data storing and requesting. For the web page design and web development, the open source toolkit ([Bootstrap 4.6](https://getbootstrap.com/)) was utilized. EnsDTI-kinase is tested on Chrome, Firefox, Microsoft Edge, and Safari across different operating systems. Predicted target kinase table is presented with the [jQuery javascript library DataTables](https://datatables.net/). Interactive PCA and projection figures are generated using [Chart.js](https://www.chartjs.org/), while mutagenesis influence map and sequential importance distribution are plotted using [D3](https://d3js.org/) library.

Providing a SMILES string of a query compound on the EnsDTI-kinase platform can quickly start searching for potential target proteins on 426 kinase proteins. On the getting-start page, users can also select one protein for residual mutagenesis significance prediction and up to ten representative proteins for probable binding sites prediction.

## 4. Results

### 4.1. Performance Evaluation

We evaluated our model using the following schemes as mentioned in Section 3.4 and illustrated in Figure 3.4:

- Internal Cross-Validation
- External Validation
- Ensemble Consensus Validation

**Figure 3:**
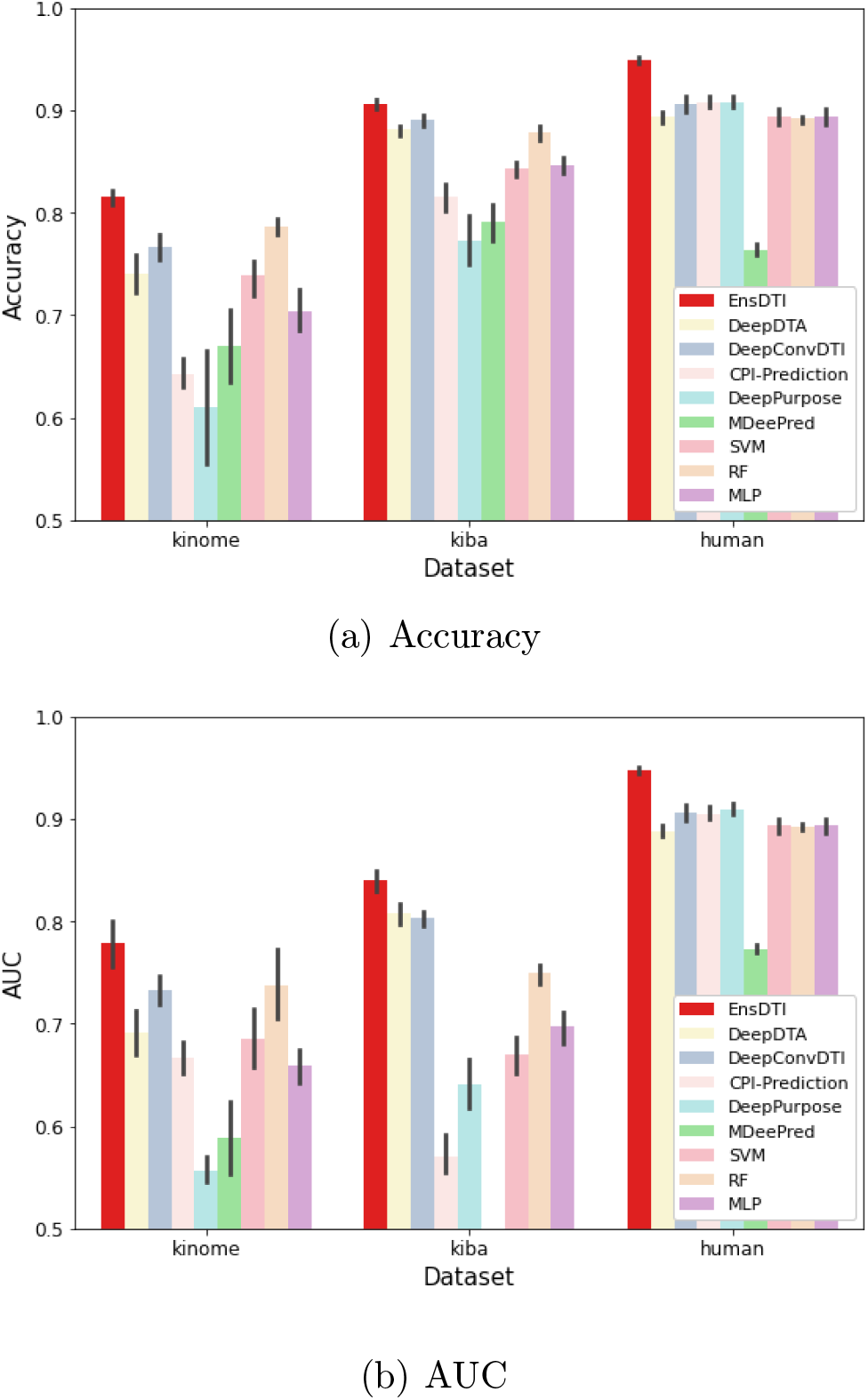
Internal validation performances under (a) Accuracy and (b) AUC metrics on all three datasets: kinome, kiba and human.

#### 4.1.1. Internal Cross-Validation

We first evaluate our model and the base classifiers under the 5-fold CV scheme on davis, kiba, kinome and human datasets (Figure 4.1.1, Table S1). Overall, EnsDTI-kinase achieved the best performance in terms of accuracy and AUC for the three datasets. On kiba data, models such as CPI-Prediction and MDeePred achieved low recall values (CPI-Prediction: 0.149 and MDeePred: 0.0), while EnsDTI-kinase achieved the best recall value (0.728). we note here that EnsDTI-kinase made balanced predictions on positive and negative labels, with the best F1-score (0.773), while the second-best performing model (DeepPurpose) achieved 0.531. AUPR of our model is the best at 0.840 while that of the runner-up model is 0.808 by DeepDTA. This means that our model is superior to other base classifiers in identifying true kinase inhibitor targets.

**Figure 4:**
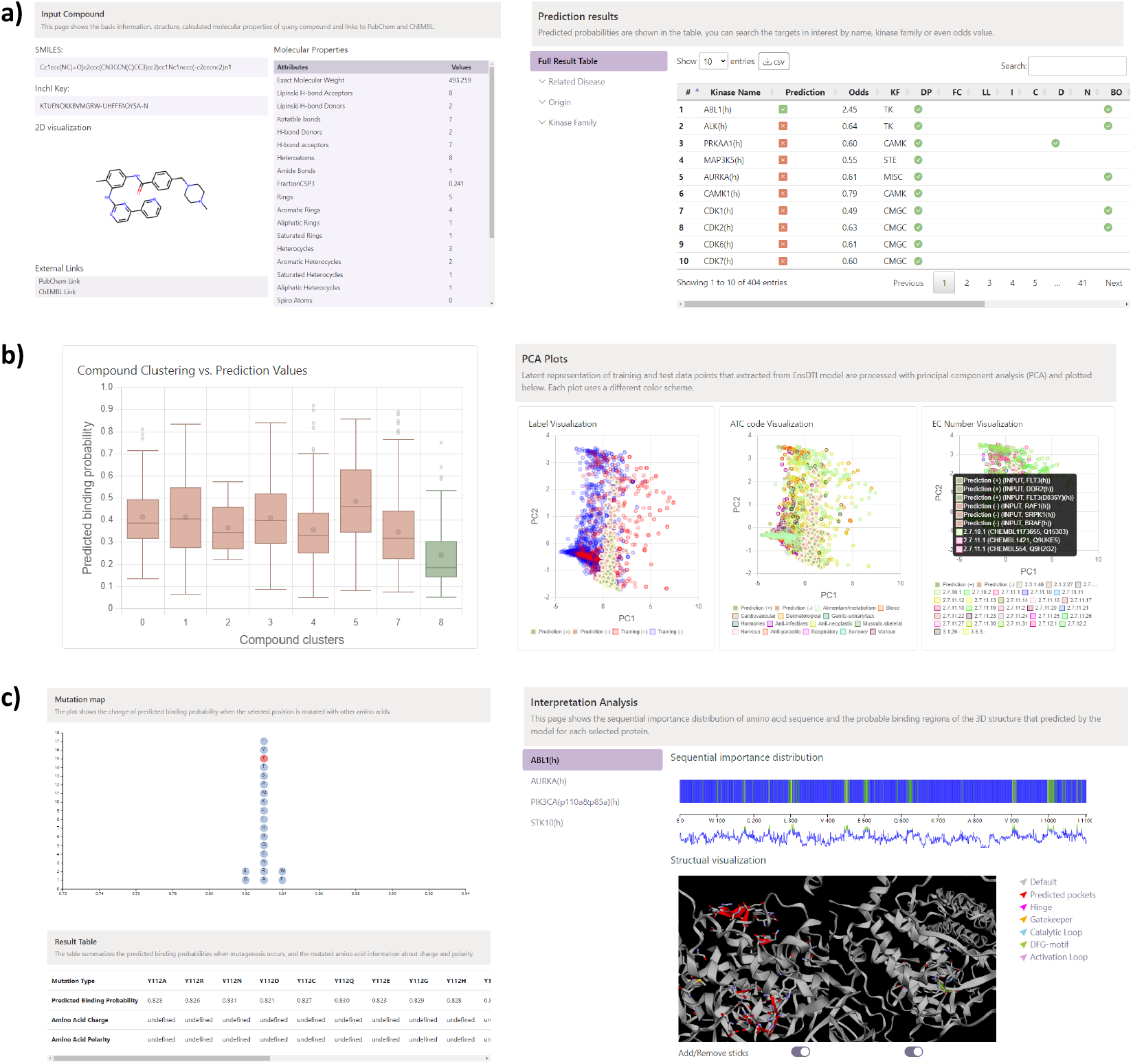
EnsDTI-kinase web-server results on case study: **a)** *In-silico* prediction on query compounds and its interaction probabilities with kinases. **b)** Exploration of query compound and predicted embedding space given known interaction relationships. **c)** *Insilico* interpretation of mutation’s influence on binding and binding site prediction.

#### 4.1.2. External Validation

We performed the external validation scheme to compare the generalization power of the models on independent data sources. We obtained the results in Table 2 that EnsDTI-kinase has more reliable predictive power for independent data sources when compared with the DL base classifiers. External validation performances of models trained on human dataset are generally poor, which is likely due to its small data size of 6,212 DTIs. Overall, EnsDTI-kinase showed stable prediction power and best-achieved performance on the external validation experiments.

**Table 2:** External validation performance of EnsDTI-kinase and base classifiers under balanced accuracy.

| Test Data | Train Data | EnsDTI-kinase | DeepDTA | DeepConv-DTI | CPI-Prediction | MDeePred | DeepPurpose |
| --- | --- | --- | --- | --- | --- | --- | --- |
| davis | human | **0.489** | 0.219 | 0.301 | 0.094 | 0.481 | 0.451 |
|  | kiba | **0.895** | 0.198 | 0.199 | 0.165 | 0.215 | 0.193 |
|  | kinome | **0.930** | 0.579 | 0.083 | 0.878 | 0.711 | 0.866 |
| human | davis | **0.833** | 0.496 | 0.467 | 0.458 | 0.458 | 0.398 |
|  | kiba | **0.761** | 0.462 | 0.493 | 0.536 | 0.521 | 0.505 |
|  | davis | **0.841** | 0.219 | 0.211 | 0.208 | 0.210 | 0.225 |
| kiba | davis | **0.877** | 0.484 | 0.525 | 0.512 | 0.478 | 0.472 |
|  | human | **0.805** | 0.708 | 0.653 | 0.763 | 0.506 | 0.539 |
|  | kinome | **0.865** | 0.387 | 0.753 | 0.294 | 0.351 | 0.285 |
| kinome | davis | **0.713** | 0.589 | 0.624 | 0.607 | 0.607 | 0.611 |
|  | human | 0.597 | **0.606** | 0.472 | 0.417 | 0.499 | 0.503 |
|  | kiba | **0.604** | 0.403 | 0.405 | 0.419 | 0.408 | 0.417 |

#### 4.1.3. Ensemble Consensus Validation

The performance of EnsDTI-kinase and DL models were compared on the same re-defined dataset (Table 3). By re-sampling the davis data, we obtained 594 and 1,912 positives and negatives in training data, while 1,699 positives and 862 negatives in test set (See Section 3.4). We observed that EnsDTI-kinase achieved better performances compared to the other state-of-the-art (SOTA) models. EnsDTI-kinase achieved 0.706 in accuracy, while the best-performed base classifier (DeepPurpose) achieved 0.538. It is also notable that combining the results from individual classifiers significantly improved recall. EnsDTI-kinase achieved 0.756 in recall, while the next best-performing model (DeepPurpose) achieved 0.394. Overall, EnsDTI-kinase produces much less false positive DTIs than the five DL models do.

**Table 3:** Performance comparison of EnsDTI-kinase with the five DL classifiers on the resampled davis dataset. Resampling of the original davis dataset produces highly confident labels as training, while the others as test sets.

| Model | Acc. | Pre. | Spe. | Rec. | Bal. Acc. | AUC | AUPR | F1-score |
| --- | --- | --- | --- | --- | --- | --- | --- | --- |
| EnsDTI-kinase | **0.706** | 0.791 | 0.607 | 0.756 | **0.682** | **0.753** | **0.856** | **0.773** |
| CPI-Prediction | 0.429 | 0.732 | 0.842 | 0.219 | 0.531 | 0.556 | 0.718 | 0.337 |
| DeepDTA | 0.404 | 0.765 | 0.912 | 0.146 | 0.529 | 0.533 | 0.704 | 0.245 |
| DeepConv-DTI | 0.454 | 0.791 | 0.875 | 0.241 | 0.558 | 0.569 | 0.729 | 0.369 |
| MDeePred | 0.342 | 0.710 | **0.990** | 0.013 | 0.501 | 0.533 | 0.684 | 0.025 |
| DeepPurpose | 0.538 | **0.813** | 0.821 | 0.394 | 0.608 | 0.570 | 0.765 | 0.531 |
| SVM | 0.342 | 0.607 | 0.022 | **0.972** | 0.497 | 0.042 | 0.497 | 0.662 |
| RF | 0.408 | 0.670 | 0.211 | 0.795 | 0.503 | 0.321 | 0.503 | 0.665 |
| MLP | 0.376 | 0.748 | 0.089 | 0.941 | 0.515 | 0.159 | 0.515 | 0.671 |

**Table 4:** Ablation study performance of EnsDTI-kinase.

| ML Models | DL Models |  |  |  | Metrics |  |  |  |  |  |  |
| --- | --- | --- | --- | --- | --- | --- | --- | --- | --- | --- | --- |
|  | (A, B) | + C | + D | + E | Accuracy | Precision | Recall | Specificity | Balanced Accuracy | AUC | AUPR |
|  | ✓ |  |  |  | 0.455 | 0.700 | 0.311 | 0.737 | 0.524 | 0.479 | 0.680 |
|  | ✓ | ✓ |  |  | 0.609 | 0.651 | 0.883 | 0.068 | 0.476 | 0.426 | 0.606 |
|  | ✓ | ✓ | ✓ |  | 0.660 | 0.663 | 0.991 | 0.009 | 0.500 | 0.565 | 0.708 |
|  | ✓ | ✓ | ✓ | ✓ | 0.651 | 0.662 | **0.968** | 0.027 | 0.497 | 0.480 | 0.669 |
| ✓ | ✓ |  |  |  | 0.546 | **0.825** | 0.401 | **0.832** | 0.617 | 0.661 | 0.799 |
| ✓ | ✓ | ✓ |  |  | 0.639 | 0.763 | 0.662 | 0.594 | 0.628 | 0.696 | 0.824 |
| ✓ | ✓ | ✓ | ✓ |  | 0.675 | 0.729 | 0.813 | 0.404 | 0.608 | 0.685 | 0.821 |
| ✓ | ✓ | ✓ | ✓ | ✓ | **0.706** | 0.791 | 0.756 | 0.607 | **0.681** | **0.753** | **0.856** |

A: CPI-Prediction, B: DeepDTA, C: DeepConv-DTI, D: MDDeePred, E: DeepPurpose

### 4.2. Ablation Analysis

EnsDTI-kinase combines predictions from eight models, thus the ablation analysis is on how many models contribute to the final prediction. Adding up each base classifier in chronological order for the ablation analysis was performed to investigate how much gain in performance our ensemble model achieved when integrating more base classifiers. Integrating more DL models as an ensemble showed balanced or better performance in different metrics. Moreover, the set of three ML models also provided 10.8% of gain on average AUC, 6.7% of gain on average balanced accuracy, and 6.0 % of gain on average AUPR to the ensemble of DL models. Overall, EnsDTI-kinase was able to make improved predictions by avoiding trends that predict all data to the same label.

### 4.3. Enrichment of Specific Data Labels

To further demonstrate our model, we investigated how much our model can better predict DTIs to specific kinase families or chemical features. To do so, PubChem fingerprints and EC number information were used to investigate the enrichment of correct DTI prediction for compounds and kinases, respectively. We found from the results that there were two kinase families (receptor and non-specific protein-tyrosine kinases) and 39 PubChem fingerprints that our model showed statistically significantly more predictive power than accuracy of 0.706 (Table 5 and Table 6).

**Table 5:** EC Numbers that show greater performance than overall EnsDTI-kinase accuracy (0.706).

| EC Number | Description | Accuracy | f1-score | p-value |
| --- | --- | --- | --- | --- |
| 2.7.10.1 | receptor protein-tyrosine kinase | 0.774 | 0.858 | 5.18e-10 |
| 2.7.10.2 | non-specific protein-tyrosine kinase | 0.729 | 0.806 | 1.29e-2 |

**Table 6:** Top 10 PubChem fingerprints that show greater performance than overall EnsDTI-kinase accuracy (0.706).

| PubChem FP ID | Description | Accuracy | f1-score | p-value |
| --- | --- | --- | --- | --- |
| 397 | N(∼C)(:C)(:C) | 0.767 | 0.772 | 2.18e-5 |
| 662 | O-C-C-O-C | 0.765 | 0.754 | 4.21e-6 |
| 567 | O-C-C-O | 0.764 | 0.754 | 5.95e-6 |
| 213 | $\geq 1$ any ring size 7 | 0.762 | 0.738 | 3.79e-5 |
| 218 | $\geq 1$ unsaturated non-aromatic nitrogen-containing ring size 7 | 0.762 | 0.738 | 3.79e-5 |
| 219 | $\geq 1$ unsaturated non-aromatic heteroatom-containing ring size 7 | 0.762 | 0.738 | 3.79e-5 |
| 737 | Cc1cc(N)ccc1 | 0.760 | 0.746 | 3.95e-5 |
| 800 | CC1CC(N)CCC1 | 0.760 | 0.746 | 3.95e-5 |
| 157 | $\geq 3$ any ring size 5 | 0.760 | 0.751 | 3.73e-5 |
| 612 | N-C-O-C-C | 0.759 | 0.741 | 1.04e-4 |
| EnsDTI-kinase |  | 0.706 | 0.773 | - |

### 4.4. Case Studies

In this section, we demonstrate our results using case studies of target prediction on kinase panel, *in silico* mutagenesis and binding site prediction. How to use EnsDTI-kinase web-server is also demonstrated by using Imatinib as a query compound.

#### 4.4.1. Target Prediction on Kinase Panel

Figure 4.4A shows ‘Target Prediction’, ‘Result Projection’, and ‘PCA Exploration’ sub-pages of EnsDTI-kinase website. ‘Target Prediction’ sub-page includes a series of bar charts (left) and a summary table (right). Each bar chart represents the predicted binding probability of a kinase group, and the summary table containing detailed information is user-interactive and supports functions like filtering, sorting, searching and downloading.

‘Result Projection’ sub-page presents two types of result visualization, compound cluster plot (left) and odds distribution plot (right), so that users can compare results with training set. Compound cluster plot shows the prediction distribution of training compound clusters and marks the belonging of query compound. Specifically, by calculating the ECFP similarity, we clustered training compounds into eight groups and mapped the query compound to the closest one. From the plot we can investigate to what extent compound structures affect prediction results. Odds distribution plot illustrates the log-odds distribution of both training data and user query. Log-odds are calculated with Equation 4. The higher the log odds, the more selective the query compound is to the specific target. ‘PCA Exploration’ sub-page visualizes model latent vectors with PCA figures. These latent vectors are obtained from the final layer of EnsDTI-kinase and implies how model makes decision on training and query data. PCA figures are colored in three schemes (interaction label, compound ATC code, and kinase EC number) so that we can infer how well the model distinguish positive and negative data, and whether different chemical characteristics and corresponding enzyme-catalyzed reaction features are captured by the model. All three PCA plots are explorable by clicking the points on the figure.

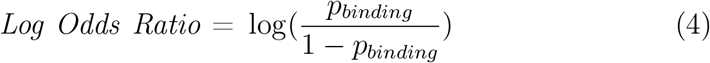

#### 4.4.2. In-silico Mutagenesis Analysis

For the kinase and specific amino acid position of interest submitted by user, EnsDTI-kinase replaces the corresponding amino acid by applying all possible amino acid residues (*in silico* point mutagenesis) and presented the results in the ‘Mutagenesis Analysis’ sub-page as shown in Figure 4.4B. The degree of perturbation in binding affinity is measured by deviations from that of the original amino acid sequence. Take the binding between Imatinib and kinase ABL1 as example, the binding affinity for ABL1(M351T) is predicted as 0.717 while the binding prediction between original ABL1 and Imatinib is 0.710. The results are in agreement with the experimental results from Apperley [40] that Imatinib is 5% - 10% much more sensitive to ABL1(M351T) than ABL1.

#### 4.4.3. Binding Site Prediction

We conducted case studies on Imatinib and four selected kinases: ABL(h), AURK(h), PIK3CA(p110a&p85a)(h) and STK10(h). The predicted binding sites are shown in Figure 5. For each amino acid sequence, top 10% of loci that are important to inhibition are marked in yellow in the heatmap. These include L300-Q306, K904-L910, and A999-G1003 for ABL(h), G10-A14, Y196-A202, A330-K338 for AURK(h), L698, R764, T1030-F1038 for PIK3CA(p110a&p85a)(h), I92-D102, S251-P253 for STK10(h). Binding site prediction results are shown in the ‘Pocket Prediction’ sub-page on the web-server.

**Figure 5:**
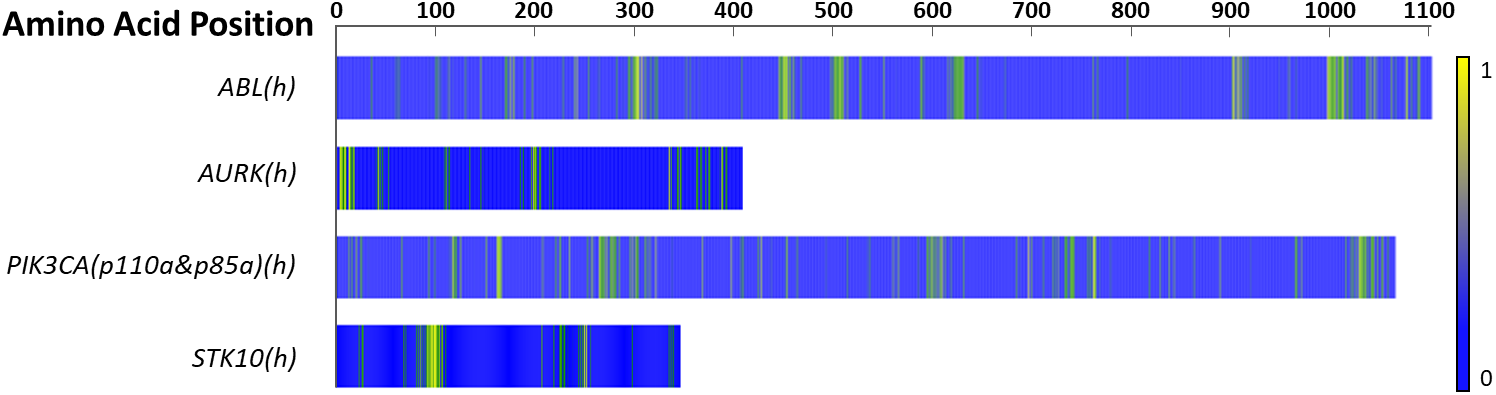
Binding site prediction heatmap for Imatinib to selected kinases: ABL(h), AURK(h), PIK3CA(p110&p85a)(h), and STK10(h). Yellow-colored regions indicate highly-likely binding sites, while blue-colored regions are less likely to be binding sites.

Our web-server also provides user-interactive 3D visualization for comparing interpretation results with KLIFS kinase motif annotation [41, 42]. Referring to the KLIFS annotation, our web-server also displays important protein residues with different colors.

Users can be informed about inhibitor types of their compounds whether they bind directly near hinge or allosteric sites. Figure 6 is the 3D visualization of the case study on KIT and Imatinib (PDB ID: 1T46). Residues that are predicted possibly for the pocket by the model are colored in red, while others are colored in gray. Imatinib is highlighted in yellow. We can see that EnsDTI-kinase gave higher weight to the locations near where binding occurs.

**Figure 6:**
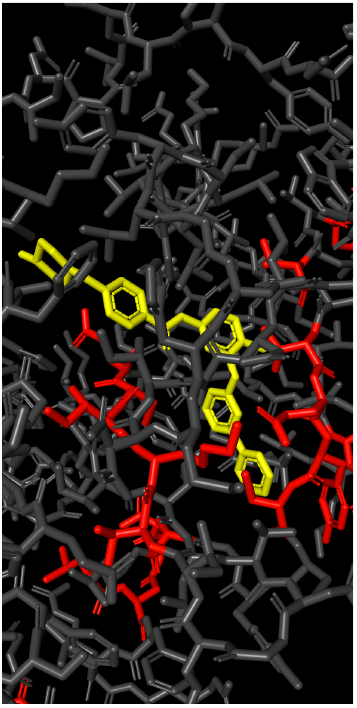
3D Visualization of pocket prediction result on KIT and Imatinib (PDB ID: 1T46). Red: Significant substructure of KIT predicted by EnsDTI-kinase. Gray: The rest of the KIT. Yellow: Imatinib.

#### 4.4.4. Analysis on Recent Drugs

In order to demonstrate our model’s performance in recent experiments, EnsDTI-kinase was evaluated on experimental results obtained from recent publications. Elkins et al. [43] tested 367 lead compounds on 224 kinases. We filtered out compounds and proteins whose sequences are already used for training, and tested the remaining DTI pairs for evaluation.

On the high-throughput profiling of available kinase inhibitors against a panel of 300 recombinant protein kinases from [5], we applied activity threshold of *pK*_*d*_ = 7 and acquired 17 cold drugs and 9,798 interactions with 404 protein kinases. Our model predicted these interactions with an overall accuracy of 0.69. On close examination of the best-predicted 8 out of 17 recent drugs, our model successfully predicted their interactions with kinases at accuracy of 0.82. We provide the complete prediction performance for 17 approved inhibitors in Supplement table S2.

## 5. Conclusion

We developed an ensemble model, EnsDTI-kinase, that integrates three machine learning and five deep learning-based methods in a single computational framework, to predict kinase-inhibitor interactions. In extensive experiments, we notice that existing DTI models tend to produce high false negative rates and have difficulty in training on databases that are imbalanced in label distribution. Significance of EnsDTI-kinase is to reduce false negatives of kinase targets while improving the overall prediction power. More importantly, EnsDTI-kinase is a computational platform where newly developed DTI tools can be easily incorporated without modifying component tools so that the DTI prediction quality can improve over time. In addition, EnsDTI-kinase provides useful functionalities for users to further investigate predicted DTI so that confidence in predicted kinase targets can be evaluated on the web. In future work, we will continue to improve EnsDTI-kinase by updating the base classifiers with state-of-the-art methods and developing more useful functions for accurate DTI prediction. In particular, we plan to incorporate protein structure information by Alphafold2 [38] and pocket information such as DUD-E [44] for kinase target prediction.

## 6. Acknowledgements

This research was supported by the Collaborative Genome Program for Fostering New Post-Genome Industry of the National Research Foundation (NRF) funded by the Ministry of Science and ICT (MSIT) (No. NRF-2014M3C9A3063541), by the Bio & Medical Technology Development Program of the National Research Foundation (NRF) & funded by the Korean government (MSIT) (No. 2022M3E5F3085677), by Institute of Information & communications Technology Planning & Evaluation (IITP) grant funded by the Korea government (MSIT) [NO. 2021-0-01343, Artificial Intelligence Graduate School Program (Seoul National University)], and by a grant (No. DY0002258224) from Ministry of food and Drug Safety in 2020.

## Supplementary Information

**Table S1:** Five-fold CV performance of EnsDTI-kinase and the eight base classifiers on the three data sets: human, kiba, and kinome.

| Model | Acc. | Pre. | Spe. | Rec. | Bal.Acc. | AUC | AUPR | F1 |
| --- | --- | --- | --- | --- | --- | --- | --- | --- |
| human |  |  |  |  |  |  |  |  |
| EnsDTI-kinase | 0.948 ± 0.0040 | 0.950 ± 0.0260 | 0.956 ± 0.0293 | 0.939 ± 0.0332 | 0.948 ± 0.0042 | 0.952 ± 0.0044 | 0.948 ± 0.0042 | 0.931 ± 0.0137 |
| DeepDTA | 0.893 ± 0.0062 | 0.865 ± 0.0103 | 0.951 ± 0.0020 | 0.825 ± 0.0125 | 0.888 ± 0.0063 | 0.906 ± 0.0057 | 0.888 ± 0.0063 | 0.849 ± 0.0101 |
| DeepConvDTI | 0.906 ± 0.0093 | 0.919 ± 0.0130 | 0.906 ± 0.0128 | 0.905 ± 0.0182 | 0.905 ± 0.0094 | 0.912 ± 0.0080 | 0.905 ± 0.0094 | 0.883 ± 0.0111 |
| CPI-Prediction | 0.907 ± 0.0061 | 0.901 ± 0.0068 | 0.931 ± 0.0045 | 0.879 ± 0.0107 | 0.905 ± 0.0065 | 0.916 ± 0.0047 | 0.905 ± 0.0065 | 0.876 ± 0.0069 |
| DeepPurpose | 0.908 ± 0.0068 | 0.937 ± 0.0077 | 0.890 ± 0.0120 | 0.929 ± 0.0081 | 0.910 ± 0.0064 | 0.913 ± 0.0064 | 0.910 ± 0.0064 | 0.893 ± 0.0075 |
| MDeePred | 0.763 ± 0.0064 | 0.877 ± 0.0053 | 0.653 ± 0.0142 | 0.892 ± 0.0046 | 0.773 ± 0.0055 | 0.749 ± 0.0090 | 0.773 ± 0.0055 | 0.761 ± 0.0072 |
| SVM | 0.894 ± 0.0093 | 0.903 ± 0.0124 | 0.901 ± 0.0158 | 0.886 ± 0.0169 | 0.893 ± 0.0092 | 0.902 ± 0.0089 | 0.893 ± 0.0092 | 0.867 ± 0.0115 |
| RF | 0.892 ± 0.0042 | 0.917 ± 0.0068 | 0.879 ± 0.0064 | 0.907 ± 0.0072 | 0.893 ± 0.0043 | 0.898 ± 0.0047 | 0.893 ± 0.0043 | 0.872 ± 0.0071 |
| MLP | 0.894 ± 0.0086 | 0.904 ± 0.0045 | 0.899 ± 0.0190 | 0.888 ± 0.0093 | 0.893 ± 0.0078 | 0.901 ± 0.0088 | 0.893 ± 0.0078 | 0.868 ± 0.0061 |
| kiba |  |  |  |  |  |  |  |  |
| EnsDTI-kinase | 0.906 ± 0.0051 | 0.930 ± 0.0025 | 0.952 ± 0.0074 | 0.728 ± 0.0289 | 0.840 ± 0.0116 | 0.941 ± 0.0048 | 0.840 ± 0.0116 | 0.923 ± 0.0043 |
| DeepDTA | 0.881 ± 0.0056 | 0.918 ± 0.0047 | 0.932 ± 0.0059 | 0.684 ± 0.0270 | 0.808 ± 0.0118 | 0.925 ± 0.0052 | 0.808 ± 0.0118 | 0.909 ± 0.0064 |
| DeepConvDTI | 0.890 ± 0.0063 | 0.913 ± 0.0059 | 0.951 ± 0.0054 | 0.656 ± 0.0213 | 0.804 ± 0.0083 | 0.932 ± 0.0056 | 0.804 ± 0.0083 | 0.907 ± 0.0072 |
| CPI-Prediction | 0.816 ± 0.0145 | 0.815 ± 0.0148 | 0.992 ± 0.0024 | 0.149 ± 0.0455 | 0.570 ± 0.0222 | 0.895 ± 0.0098 | 0.570 ± 0.0222 | 0.815 ± 0.0149 |
| DeepPurpose | 0.773 ± 0.0286 | 0.847 ± 0.0255 | 0.869 ± 0.0178 | 0.414 ± 0.0444 | 0.641 ± 0.0286 | 0.858 ± 0.0200 | 0.641 ± 0.0286 | 0.840 ± 0.0254 |
| MDeePred | 0.790 ± 0.0205 | 0.790 ± 0.0205 | 1.000 ± 0.0000 | 0.000 ± 0.0000 | 0.500 ± 0.0000 | 0.883 ± 0.0128 | 0.500 ± 0.0000 | 0.790 ± 0.0205 |
| SVM | 0.843 ± 0.0078 | 0.854 ± 0.0078 | 0.965 ± 0.0068 | 0.376 ± 0.0447 | 0.670 ± 0.0202 | 0.906 ± 0.0068 | 0.670 ± 0.0202 | 0.852 ± 0.0085 |
| RF | 0.878 ± 0.0080 | 0.886 ± 0.0079 | 0.969 ± 0.0071 | 0.529 ± 0.0285 | 0.749 ± 0.0116 | 0.926 ± 0.0067 | 0.749 ± 0.0116 | 0.883 ± 0.0087 |
| MLP | 0.847 ± 0.0087 | 0.866 ± 0.0079 | 0.953 ± 0.0068 | 0.440 ± 0.0414 | 0.697 ± 0.0183 | 0.908 ± 0.0074 | 0.697 ± 0.0183 | 0.862 ± 0.0089 |
| kinome |  |  |  |  |  |  |  |  |
| EnsDTI-kinase | 0.815 ± 0.0079 | 0.777 ± 0.0325 | 0.695 ± 0.1092 | 0.863 ± 0.0600 | 0.779 ± 0.0258 | 0.731 ± 0.0750 | 0.779 ± 0.0258 | 0.652 ± 0.0950 |
| DeepDTA | 0.740 ± 0.0213 | 0.692 ± 0.0709 | 0.534 ± 0.0974 | 0.848 ± 0.0464 | 0.691 ± 0.0260 | 0.602 ± 0.0890 | 0.691 ± 0.0260 | 0.549 ± 0.1162 |
| DeepConvDTI | 0.767 ± 0.0148 | 0.707 ± 0.0680 | 0.632 ± 0.0760 | 0.833 ± 0.0443 | 0.732 ± 0.0166 | 0.667 ± 0.0720 | 0.732 ± 0.0166 | 0.588 ± 0.1048 |
| CPI-Prediction | 0.643 ± 0.0163 | 0.512 ± 0.0871 | 0.861 ± 0.0640 | 0.473 ± 0.0982 | 0.667 ± 0.0175 | 0.641 ± 0.0859 | 0.667 ± 0.0175 | 0.494 ± 0.0971 |
| DeepPurpose | 0.610 ± 0.0646 | 0.496 ± 0.0885 | 0.328 ± 0.0211 | 0.786 ± 0.0307 | 0.557 ± 0.0144 | 0.393 ± 0.0384 | 0.557 ± 0.0144 | 0.427 ± 0.1025 |
| MDeePred | 0.670 ± 0.0412 | 0.597 ± 0.1043 | 0.309 ± 0.1403 | 0.868 ± 0.0638 | 0.589 ± 0.0395 | 0.400 ± 0.1448 | 0.589 ± 0.0395 | 0.456 ± 0.1335 |
| SVM | 0.738 ± 0.0180 | 0.677 ± 0.0651 | 0.539 ± 0.1368 | 0.830 ± 0.0720 | 0.685 ± 0.0333 | 0.597 ± 0.1095 | 0.685 ± 0.0333 | 0.540 ± 0.1210 |
| RF | 0.787 ± 0.0090 | 0.760 ± 0.0447 | 0.604 ± 0.1365 | 0.871 ± 0.0584 | 0.737 ± 0.0400 | 0.668 ± 0.1017 | 0.737 ± 0.0400 | 0.606 ± 0.1165 |
| MLP | 0.704 ± 0.0228 | 0.611 ± 0.0808 | 0.538 ± 0.1115 | 0.779 ± 0.0724 | 0.658 ± 0.0204 | 0.572 ± 0.0981 | 0.658 ± 0.0204 | 0.508 ± 0.1165 |

Table S1 shows the internal validation performances of EnsDTI-kinase and all base classifiers.

Figure S1 shows the external validation performances of EnsDTI-kinase and five DL-based base classifiers. Our model shows the best and the most stable performances under different metrics (e.g. accuracy, balanced accuracy, F1, AUC).

Table S2 shows the prediction performance on recent 17 approved inhibitors. To evaluate our model on real drug data, we filtered out the compounds and proteins whose sequences are already used in the training process, and test the remaining DTI pairs from Elkins et al. [43].

Table S3 shows the confusion matrix of ensemble consensus validation experiment. Not only well predict most of the binding interactions, our model also presents the least false negative number which shows its ability to rescue potential targets for further experimental procedures.

**Figure S1:**
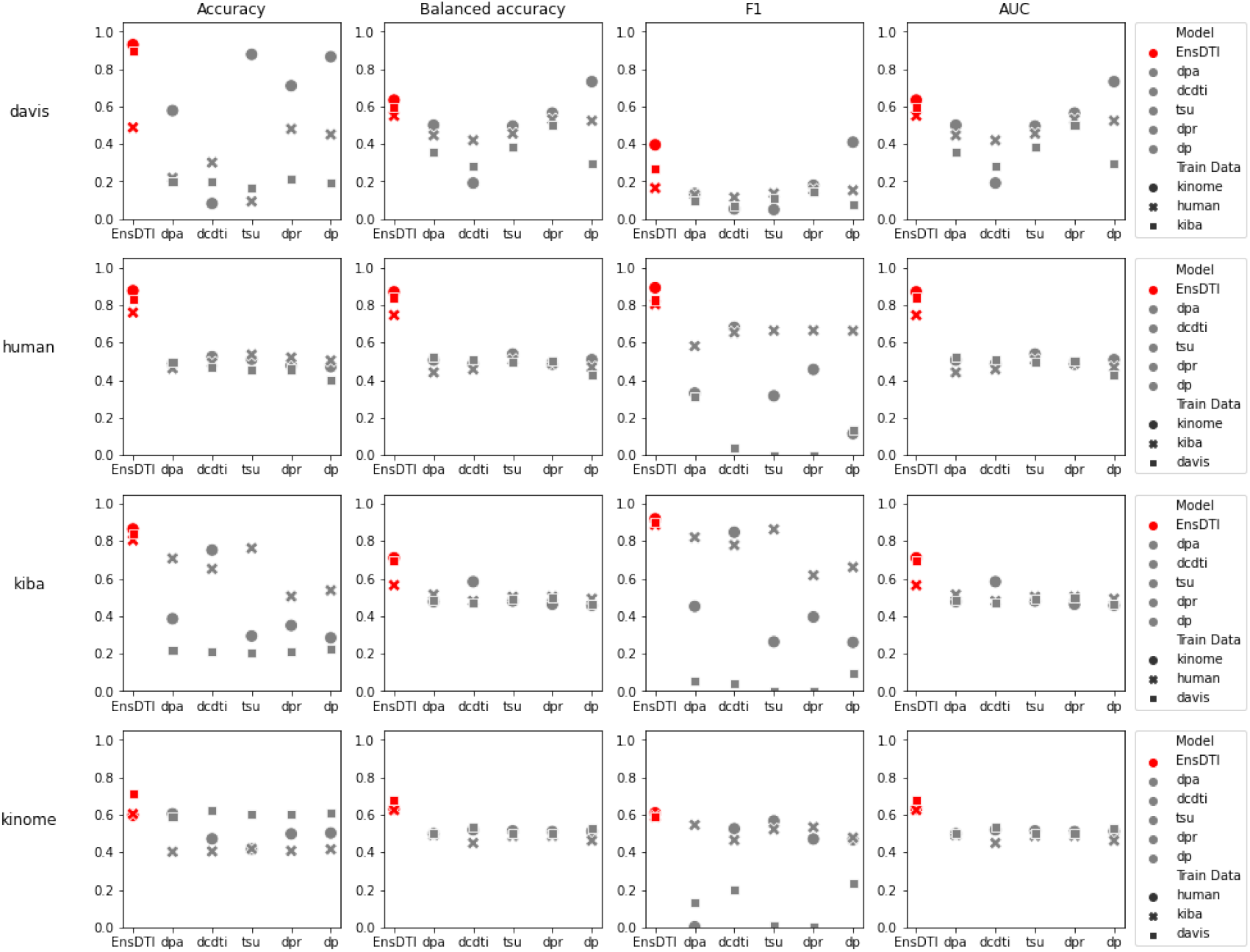
Comparison of external validation performance

**Table S2:**
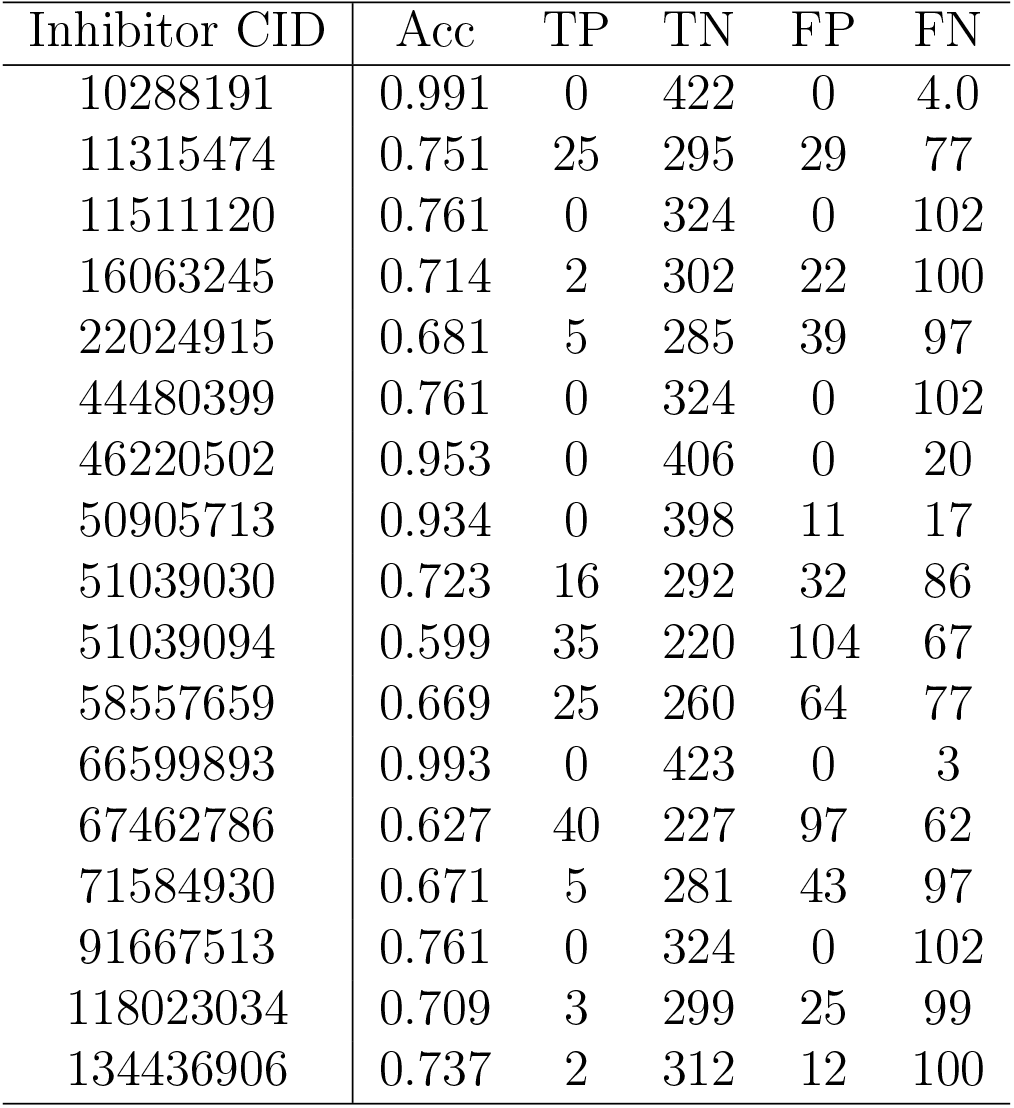
Prediction performance of EnsDTI-kinase on 17 approved kinase inhibitors (CIDs: 10288191, 11315474, 11511120, 16063245, 22024915, 44480399, 46220502, 50905713, 51039030, 51039094, 58557659, 66599893, 67462786, 71584930, 91667513, 118023034, 134436906).

| Inhibitor CID | Acc | TP | TN | FP | FN |
| --- | --- | --- | --- | --- | --- |
| 10288191 | 0.991 | 0 | 422 | 0 | 4.0 |
| 11315474 | 0.751 | 25 | 295 | 29 | 77 |
| 11511120 | 0.761 | 0 | 324 | 0 | 102 |
| 16063245 | 0.714 | 2 | 302 | 22 | 100 |
| 22024915 | 0.681 | 5 | 285 | 39 | 97 |
| 44480399 | 0.761 | 0 | 324 | 0 | 102 |
| 46220502 | 0.953 | 0 | 406 | 0 | 20 |
| 50905713 | 0.934 | 0 | 398 | 11 | 17 |
| 51039030 | 0.723 | 16 | 292 | 32 | 86 |
| 51039094 | 0.599 | 35 | 220 | 104 | 67 |
| 58557659 | 0.669 | 25 | 260 | 64 | 77 |
| 66599893 | 0.993 | 0 | 423 | 0 | 3 |
| 67462786 | 0.627 | 40 | 227 | 97 | 62 |
| 71584930 | 0.671 | 5 | 281 | 43 | 97 |
| 91667513 | 0.761 | 0 | 324 | 0 | 102 |
| 118023034 | 0.709 | 3 | 299 | 25 | 99 |
| 134436906 | 0.737 | 2 | 312 | 12 | 100 |

**Table S3:**
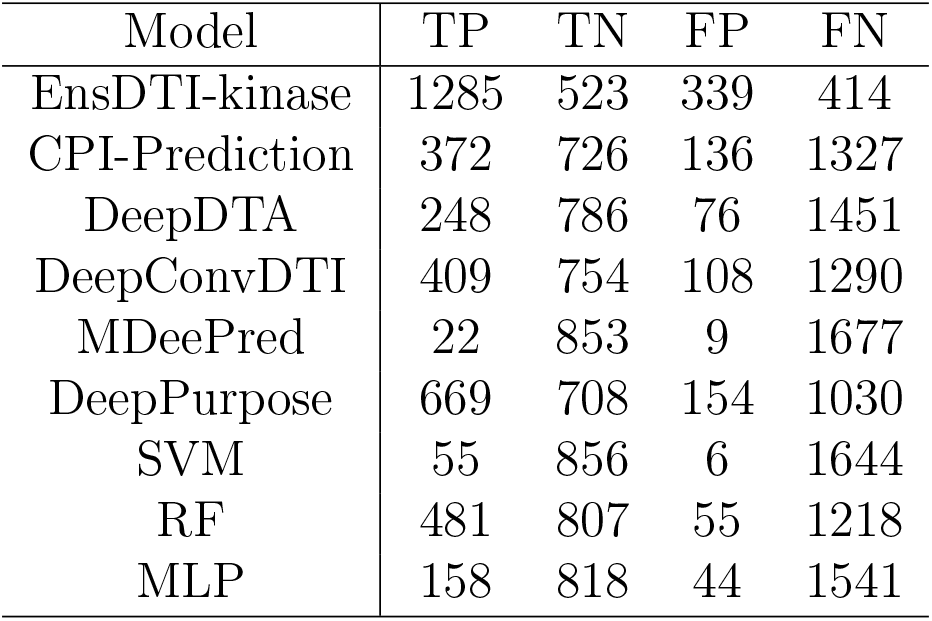
Confusion matrix of ensemble consensus validation experiment.

| Model | TP | TN | FP | FN |
| --- | --- | --- | --- | --- |
| EnsDTI-kinase | 1285 | 523 | 339 | 414 |
| CPI-Prediction | 372 | 726 | 136 | 1327 |
| DeepDTA | 248 | 786 | 76 | 1451 |
| DeepConvDTI | 409 | 754 | 108 | 1290 |
| MDeePred | 22 | 853 | 9 | 1677 |
| DeepPurpose | 669 | 708 | 154 | 1030 |
| SVM | 55 | 856 | 6 | 1644 |
| RF | 481 | 807 | 55 | 1218 |
| MLP | 158 | 818 | 44 | 1541 |

